# MetalinksDB: a flexible and contextualizable resource of metabolite-protein interactions

**DOI:** 10.1101/2023.12.30.573715

**Authors:** Elias Farr, Daniel Dimitrov, Denes Turei, Christina Schmidt, Sebastian Lobentanzer, Aurelien Dugourd, Julio Saez-Rodriguez

## Abstract

Interactions between proteins and metabolites are key for cellular function, from the catalytic breakdown of nutrients to signaling. An important case is cell-cell communication, where cellular metabolites are secreted into the microenvironment and initiate a signaling cascade by binding to an intra- or extracellular receptor of another cell. While protein-protein mediated cell-cell communication is routinely inferred from transcriptomic data, for metabolite-protein interactions this is challenging due to the limitations of high-throughput single-cell and spatial metabolomics technologies, together with the absence of comprehensive prior knowledge resources that include metabolites. Here we report MetalinksDB, a comprehensive and flexible database of intercellular metabolite-protein interactions that is a magnitude larger than existing ones. MetalinksDB can be tailored to specific biological contexts such as diseases, pathways, or tissue/cellular locations by querying subsets of interactions using the web interface (https://metalinks.omnipathdb.org/) or the knowledge graph adapters. We showcase the use of MetalinksDB by identifying deregulated processes in renal cancer patients from multi-omics data as well as inferring metabolite-mediated cell-cell communication events driving acute kidney injury from spatial transcriptomic data. We anticipate that MetalinksDB will facilitate the study of metabolite-mediated communication processes.

**Graphical Abstract:** 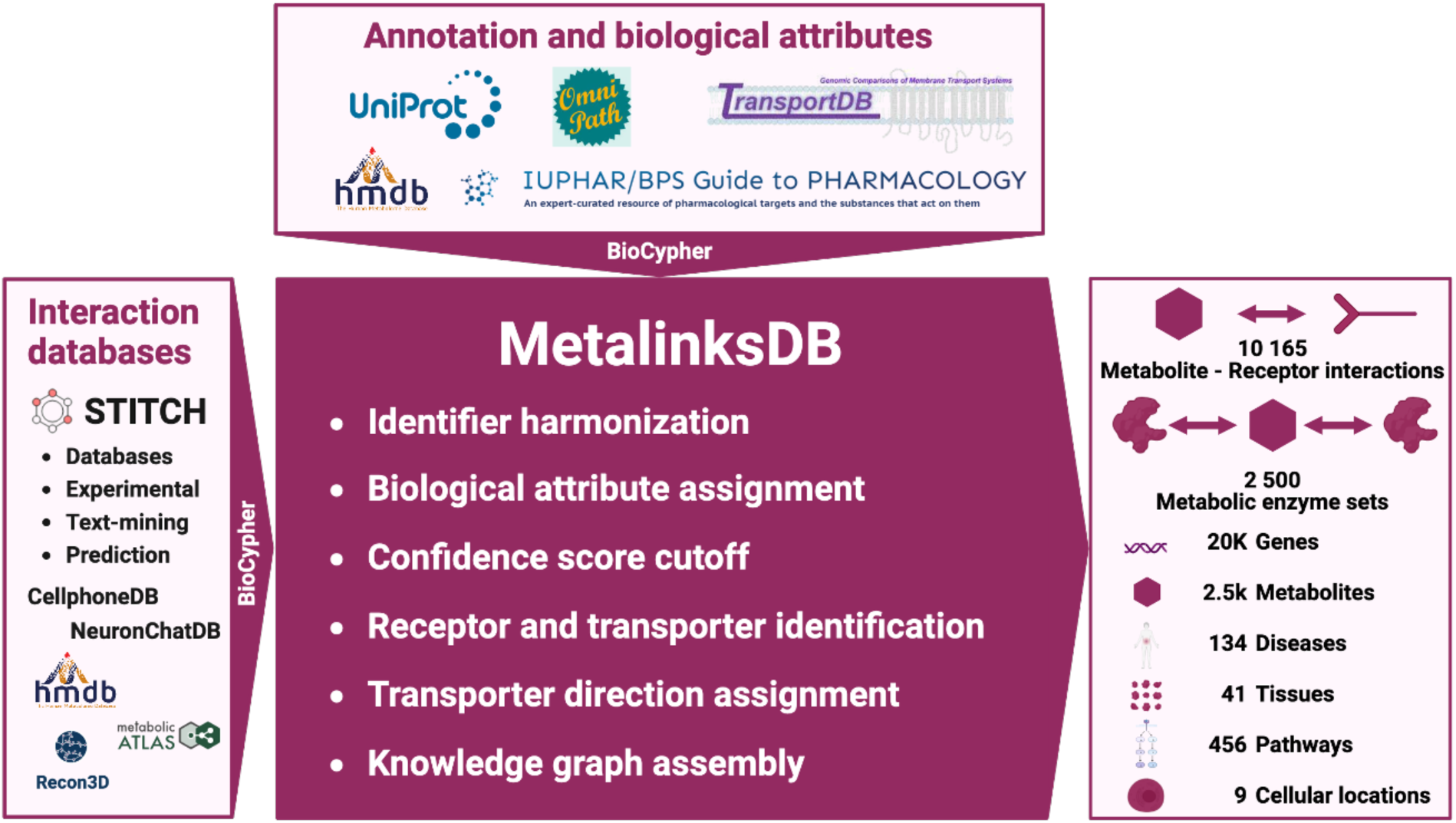

## 1. Introduction

Protein-metabolite interactions are at the center of many cellular functions. The enzymatic catalysis of metabolites and hence the use of nutrients is crucial for cellular survival, and metabolites also act as signaling molecules by regulating enzymatic activity, energy homeostasis, and signaling ^1,2^. The latter includes not only intracellular but also intercellular cell-cell communication (CCC), mediated by messenger molecules. In recent years, technological advances in high-throughput sequencing such as single-cell and spatial transcriptomics, have made the computational inference of protein-mediated CCC a standard practice^3^. The most common approach, which is largely limited to the co-expression of gene transcripts, relies on the co-expression between protein-coding genes^1^, and their contextualisation in the context of CCC via prior knowledge (prior knowledge)^4^. As such, extensive effort has been dedicated to curating ^5–7^ and gathering CCC resources^8^ focusing on protein-mediated CCC. Yet this neglects a large portion of CCC interactions, such as the binding of extracellular metabolites to receptor proteins ^2,9–12^.

Recent work has attempted to bridge this gap by gathering knowledge in the context of metabolite-mediated CCC. NeuronChat^13^ - a resource focused on the brain - and CellPhoneDBv4^14^ - a predominantly protein-protein interaction resource, originally developed for the biology of the reproductive niche^15^ - were both assembled through careful manual curation (Supplementary Table 1). Other resources such as Cellinker^16^ and scConnect^17^ get their interactions mostly from the “IUPHAR Guide to Pharmacology”^18^ while MEBOCOST^19^ collected interactions through databases like the human metabolome database (HMDB)^20^ and text-mining. Common sources for metabolite-protein interactions are the STITCH database^21^, which consists of over 20 000 000 small-molecule protein interactions from various sources, the “IUPHAR Guide to Pharmacology”^18^ and HMDB^20^. Moreover, information on metabolic enzymes, transporters, and their substrates can be found in genome-scale metabolic models (GEMs) like Recon3D^22^ or the human metabolic atlas^23^.

Despite these efforts to generate prior knowledge for metabolite-mediated CCC, the current resources are limited in coverage, focusing on specific biological niches, a set of metabolites derived from other databases, or relying heavily on manual curation (Supplementary Table 1). As such, these resources are of limited coverage, typically lack transparency in their assembly process, and tend to remain outdated to newer versions of the underlying databases.

By assembling a knowledge graph using flexible BioCypher ^24^ adapters, we integrated all of the above resources into a comprehensive, open-source, and extendable database - MetalinksDB, which is an order of magnitude larger than existing ones. In addition, MetalinksDB offers biological information about pathways, diseases, tissues etc., to allow the resource to be contextualized by removing interactions that are not relevant to a specific biological question of interest. To enable users to make contextualization queries themselves, we assembled an interactive webpage (https://metalinks.omnipathdb.org/). We demonstrate the use of MetalinksDB to analyze metabolite-protein interactions on a multi-omic data set of clear cell Renal Cell Carcinoma (ccRCC)^25^. Moreover, we use MetalinksDB with LIANA+, an all-in-one framework for CCC inference^26^, and showcase this application on a spatial transcriptomics dataset of acute kidney injury (AKI).

## 2. Results

### 2.1 MetalinksDB: transparent and reproducible integration of interaction resources

The prior knowledge currently available for metabolite-mediated CCC is limited in terms of size, reproducibility, extensibility and lacks the possibility of customization. To address these limitations, we assembled a knowledge graph taking advantage of BioCypher, a knowledge graph assembly framework that allows the straightforward incorporation of new resources and updates of existing ones in case of changes^27^.

By writing BioCypher adapters - short pieces of code that reproducibly format an input dataset - we integrated several prior knowledge resources including STITCH^21^, HMDB^20^, Recon3D^22^, and the human metabolic atlas ^23^ (see Supplementary Note 1,2). After several filtering steps and leveraging multiple annotation databases (Figure S2, Methods), we were able to obtain a high-quality knowledge graph comprising approx. 11,000 metabolite-receptor interactions as well as the metabolic enzyme sets for around 2,500 metabolites (Figure 1A, Methods). Moreover, we added biological descriptors such as diseases, pathways, and tissue locations to the nodes (metabolites and proteins) and edges (interactions) of the knowledge graph, allowing the contextualization to specific biological questions.

**Figure 1:**
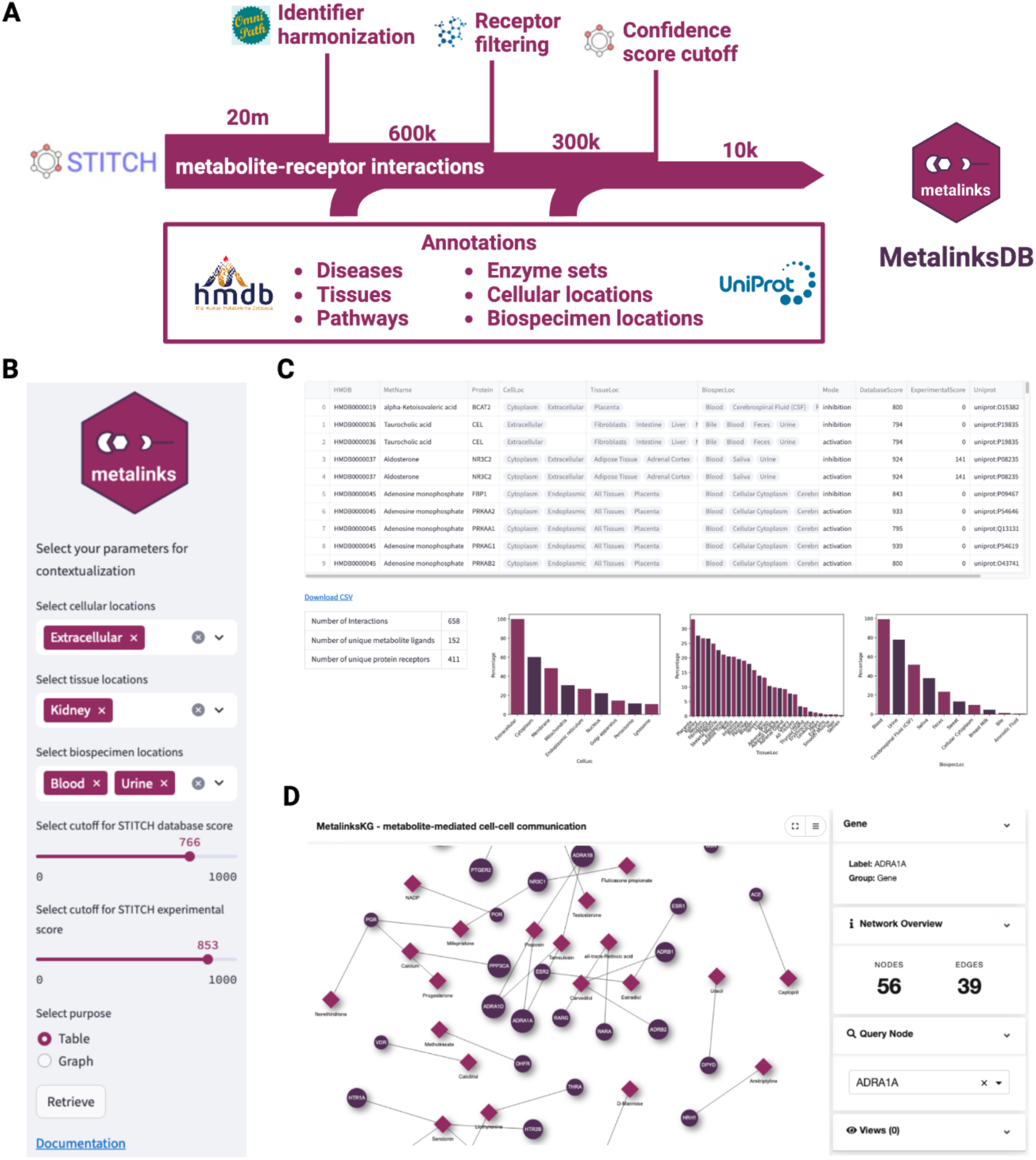
MetalinksDB graph assembly. (A) Filtering and annotation procedure during graph assembly. Over 20 million chemical-protein interactions from STITCH were filtered to the correct identifiers, receptor identity, and confidence. Moreover, annotations from several databases were added to the remaining metabolite-protein interactions. (B) Input panel for the web interface: the database can be queried for metabolites annotated to be present in a certain cellular location, tissue, or biospecimen. The bottom panel enables the cutoff for the STITCH database and experimental value to be chosen, as well as the desired output. (C) Contextualization table output panel: the contextualized table can be displayed, while in the lower panel, several control plots are shown. (D) Graph investigation output panel: a graph is visualized with metabolites as diamonds and proteins as circles through the drugst.one ^28^ interface, enabling investigation of hubs and specific interactions.

To simplify the contextualization process, we deployed a web interface available at https://metalinks.omnipathdb.org (Figure 1 B-D). This interface has two main functionalities: I) the contextualization of the MetalinksDB knowledge graph through interactive queries and II) the investigation of interactions of specific metabolites or proteins of interest. Both functionalities are accessed through a side panel that allows the input of several biological parameters such as tissue, cellular location, and biospecimen (Figure 1B). The output table and several control metrics are displayed in the main panel, a download button allows the query results to be saved as .csv (Figure 1C). On a second page, specific metabolites or proteins can be investigated by visualizing the query results as interactive graphs ^28^ (Figure 1D).

Since the annotations of metabolites can be sparse, we wanted to investigate whether the coverage of biological descriptors is high enough to enable the analysis with prior knowledge contextualized to specific conditions. We therefore quantified how many metabolites are annotated to appear in a certain disease, tissue, cellular location, or pathway. As can be seen in Figure S3, most annotations lie in the range of 2 to 20 percent, demonstrating that even with contextualized prior knowledge, the remaining coverage contains sufficient interactions.

### 2.2 MetalinksDB - a comprehensive and customizable knowledge graph

To assess the effectiveness and comprehensiveness of our database, we conducted a comparative analysis with existing databases (see Table 1). This comparison was carried out by dividing the interactions into two categories: metabolite-receptor interactions and metabolic enzyme sets (sets that allow the estimation of metabolite abundance).

For the metabolite-receptor interactions, we quantified the number of connections, ligands, and receptors involved (Figure 2A). Our analysis revealed that MetalinksDB with 11.134 interactions is an order of magnitude greater than other databases (< 650 interactions). To gain insight into the composition of these databases, we further investigated whether there are specific classes of ligands that are more prevalent in different databases (Figure 2B, S4). We could observe that MetalinksDB has a higher fraction of steroids (0.32 to 0.09, 0.18, 0.12) while having similar levels of fatty acyls (0.16 to 0.11, 0.24, 0.25) in comparison with MEBOCOST, scConnect, and Cellinker. It however has smaller fractions of carboxylic acids (0.06 to 0.19, 0.12, 0.21) (Figure 2B). Noteworthy, the database sizes differ by an order of magnitude, suggesting that MetalinksDB still has a similar content of carboxylic acids, underpinning the comprehensiveness of MetalinksDB (Figure S4D).

**Figure 2:**
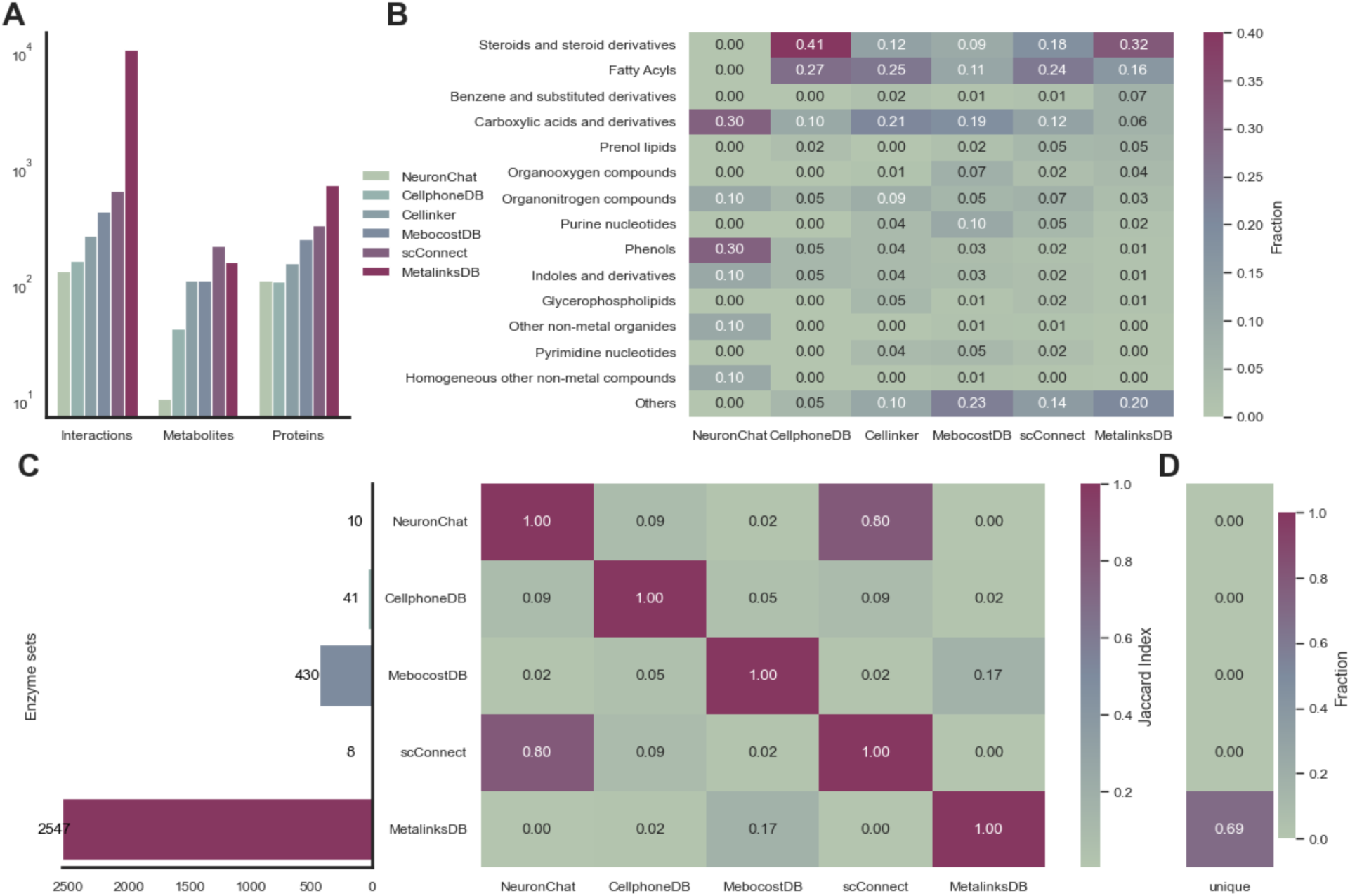
Comparison of MetalinksDB to other databases. (A) Comparison of metabolite-receptor resources. MetalinksDB contains a magnitude more interactions and the most proteins displayed, while scConnect has the most metabolites. (B) Comparison of metabolite classes showing that while MetalinksDB has the highest fraction of steroids MEBOCOST, scConnect, and Cellinker have a higher fraction of carboxylic acids. (C) Comparison of metabolic enzyme set size and overlap of enzyme sets. MetalinksDB has by far the highest number of metabolic enzyme sets associated with each metabolite. The Jaccard index heatmap shows that there is a high overlap between scConnect and NeuronChat. The highest overlap of MetalinksDB is with MebocostDB (0.17) (D) Fraction of unique sets between the databases, highlighting that only MetalinksDB contains unique metabolite sets (0.69).

Next, we compared the sizes of the metabolic enzyme sets available in different databases. Through this comparison, we observed that MetalinksDB encompasses over five times as many metabolic enzyme sets as other databases (Figure 2C). Furthermore, we assessed the overlap of metabolites that can be inferred between MetalinksDB and other databases. We found that all metabolic enzyme sets present in other databases were also contained within MetalinksDB (Figure 2C). This expansion in enzyme coverage further strengthens the comprehensiveness of MetalinksDB as a resource for estimating metabolite abundance.

### 2.3 MetalinksDB enables prior knowledge contextualization to specific disease contexts

To showcase the extended coverage of MetalinksDB and the possibility of contextualization, we examined the database’s capacity to identify metabolite-mediated CCC events and reproduce findings from the literature. Metabolite-mediated CCC plays an important role in several kidney diseases including kidney cancer which can be seen as a metabolic disease^29^. Hence, we used MetalinksDB to analyze a combined metabolomic and transcriptomics dataset of clear cell Renal Cell Carcinoma (ccRCC), a specific form of kidney cancer. To decrease the number of putative metabolite-receptor interactions, and hence potential false positives, we filtered the MetalinksDB resource to metabolites that are annotated as present in the kidney, all tissues, or the urine in HMDB and known to be extracellular (Figure 3A). This contextualization ability of MetalinksDB reduced the metabolite-receptor resource to 600 metabolite-protein interactions relevant to our specific context of kidney cancer.

**Figure 3:**
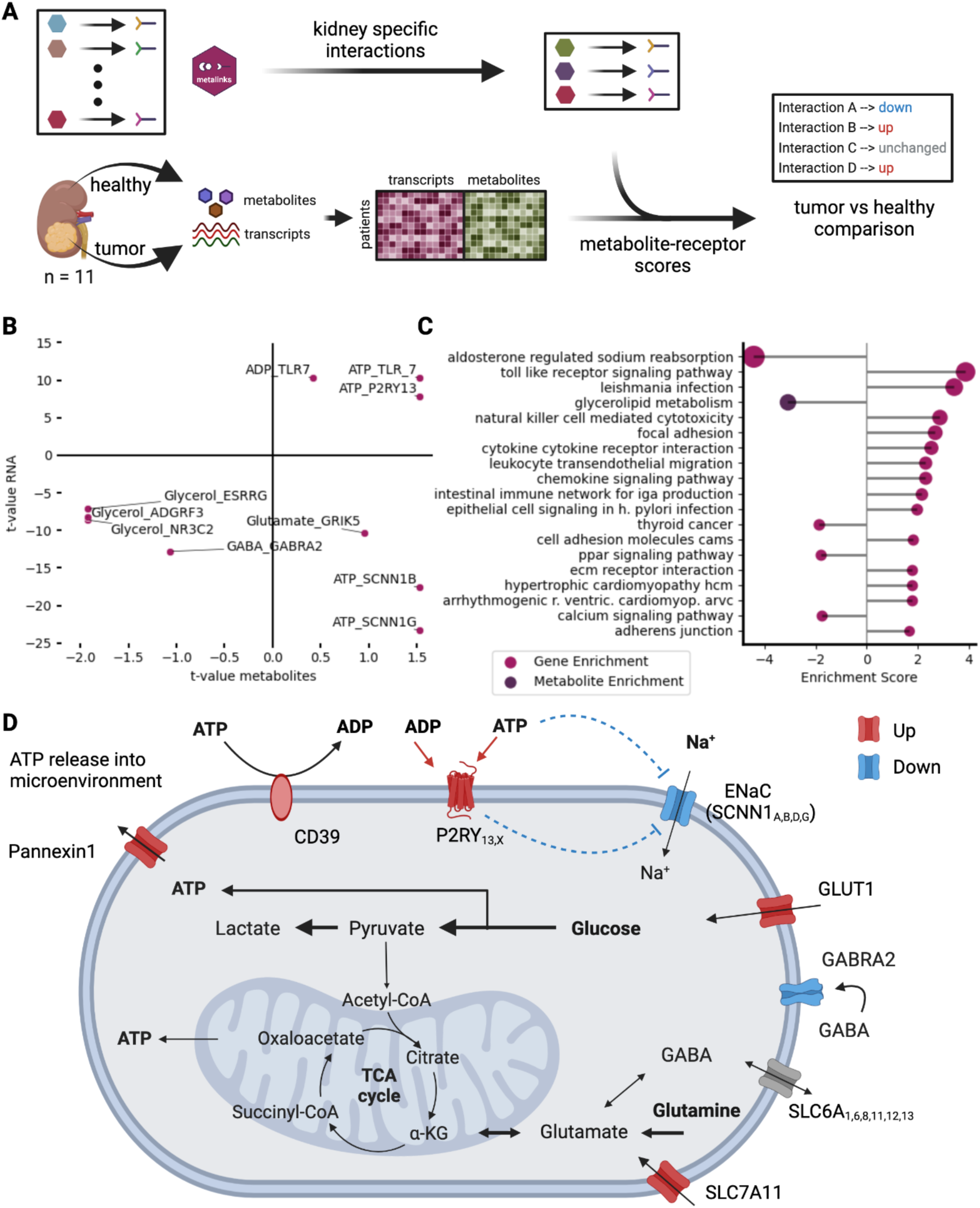
Bulk case study with contextualization. (A) Experimental strategy to infer cross-condition communication analysis using transcriptomic and metabolomic data of renal cancer patients. MetalinksDB was contextualized to metabolites known to be present extracellularly in the kidney, blood, or urine. Differential abundance analysis was performed on each omics modality independently and the t-values of a connection from MetalinksDB were averaged to get the communication score (B) Top 10 deregulated interactions shown by the t-value of the transcriptome and metabolome differential abundance analysis. While in the upper right quadrant, many ATP and ADP interactions are displayed, the lower right quadrant shows interactions with ATP and ENac subunits (SCNN1_B,G_) (C) Deregulated pathways of metabolite and transcript pathway enrichment using univariate linear models. The highest enriched transcriptome interaction is “aldosterone-regulated sodium reabsorption”, while the highest enriched metabolite pathway is glycerolipid metabolism.

The ccRCC dataset consists of metabolic and transcriptomic analysis of 11 kidney cancer patients comparing tumor versus healthy tissue ^25^ (Figure 3A). Using these ccRCC data, we calculated communication scores for putative metabolite-receptor interactions that are deregulated in tumor tissue using the mean of the t-values as input (see Methods)(Figure 3B, Supplementary Table 2). In ccRCC, an upregulation of anaerobic glycolysis is known to be important for the production of Adenosine Triphosphate (ATP) ^30,31^, which is not only an important source of energy but can also be a signaling molecule ^32–34^. In line with this, we found multiple metabolite-receptor interactions where both ATP and Adenosine Diphosphate (ADP) are involved (Fig. 3B). Interestingly, the nucleotide precursor adenosine, which is downregulated in tumors compared to healthy tissue, was connected to the Adenosine A2b Receptr (ADORA2B) in the original publication ^25^, which we also find in our Metalinks results, yet not amongst the top interactions (Supplementary Table 2). As the next step, we performed pathway enrichment analysis using KEGG pathways ^35^ and the metabolite-receptor interaction score and found “aldosterone-regulated sodium reabsorption” and “Toll-like receptor signaling pathway” among the top dysregulated pathways. Both of these pathways contain receptors that are potentially bound (TLR7), activated (TLR3), or inhibited (SCNN1B, SCNN1G) by ATP (Fig. 3C) and are known to play a role in kidney physiology ^36–38^ or ccRCC ^39^. SCNN1B and SCNN1G are both part of the epithelial sodium transporter ENaC ^40^, known markers of ccRCC ^41^, and are implicated with sodium wasting in a SCNN1B genetic condition^42^. In line with this,the pathway “aldosterone-regulated sodium reabsorption” is downregulated in the pathway analysis (Fig. 3C). Lastly, we found the interaction of ATP and the purinergic receptor P2RY13 (Fig. 3B), which is part of a receptor class known to be activated by ATP and ADP amongst other purines and pyrimidines ^43–45^. Connecting those metabolite-receptor interactions found using MetalinksDB we hypothesized that the increased levels of ATP (Log2FC 2.33, p.val 0.14) in tumor cells, likely originate from the increased anaerobic glycolysis, could be released via the transporter Pannexin 1, which was also upregulated in tumor cells (Log2FC 1.00, p.val 2.76 x 10^-4^) and in turn activate purinergic receptor whilst further inhibiting sodium transporters. The latter might in turn lead to diminished sodium levels in ccRCC, which has been previously proposed to serve as a prognostic and predictive factor of metastatic ccRCC^46^ (Fig. 3D). Together this showcases MetalinkDB’s ability to find disease-specific, coherent metabolite-receptor interactions for hypothesis generation, which can not be found with current databases (Fig S5) and can be further investigated by dedicated experiments.

### 2.4 Using MetalinksDB to infer metabolite-mediated interactions driving kidney injury

Due to the absence of comprehensive single-cell metabolomic datasets, recently developed tools infer metabolite abundance from transcriptomic data ^13,14,19^ and then infer a communication score with the known receptors of these predicted metabolite abundances; thus, generating hypotheses of putative metabolite-receptor interactions (Figure 4A). Both steps require extensive prior knowledge - available in MetalinksDB (Figure 4A).

**Figure 4:**
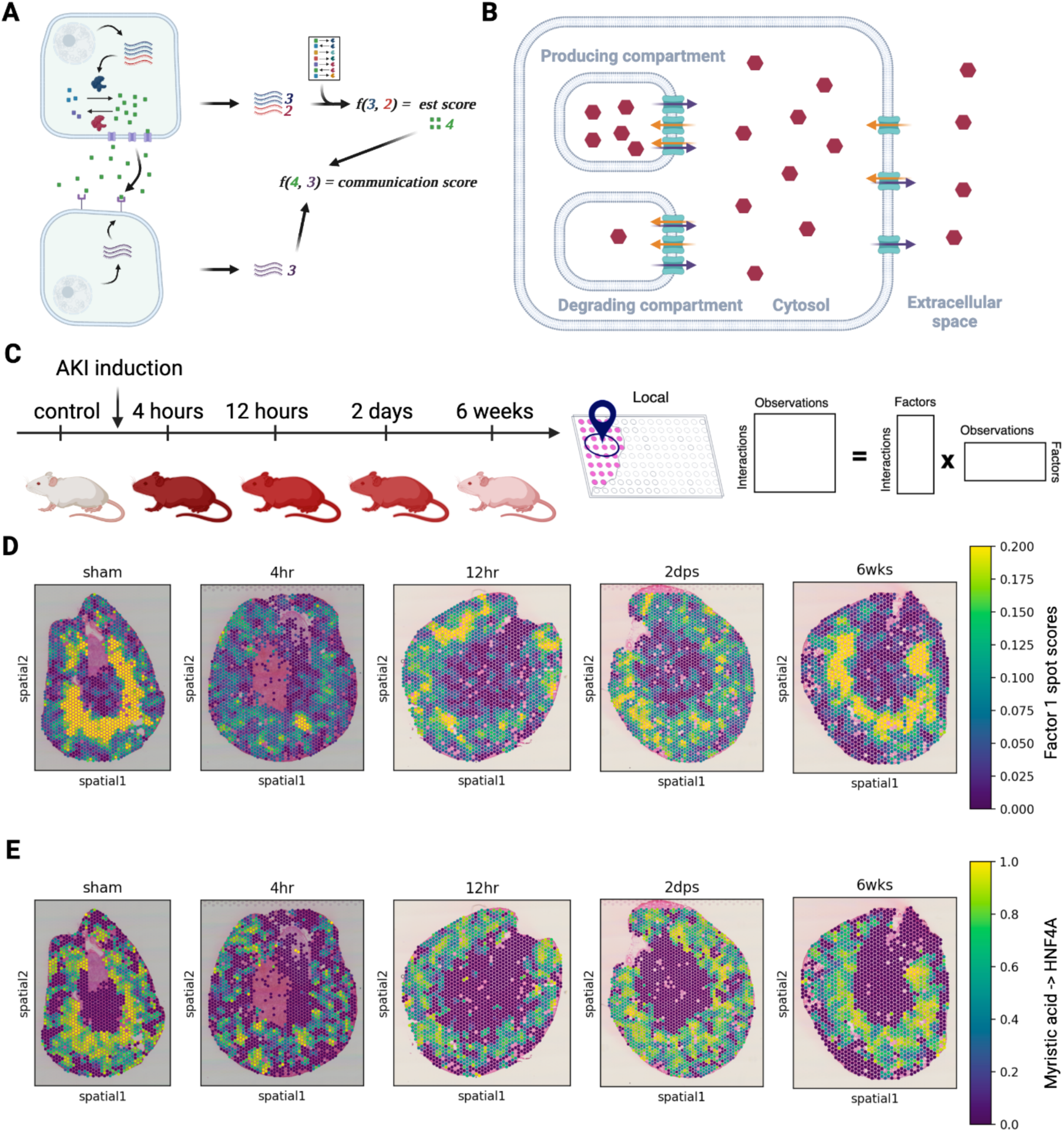
Metabolite-mediated CCC inference in AKI using LIANA+. (A) Principle of metabolite-mediated CCC inference from transcriptomics (B) Visualization of importers and exporters (C) Experimental setup to study murine kidney injury. AKI was induced in mice and samples for spatial sequencing were taken at 4 and 12 hours as well as 2 days and 6 weeks after treatment. (D) Factor 1 spot scores in mouse acute kidney injury spatial transcriptomics. The factor describes interactions that are strongly present in the control (sham), disappear during injury and increase to sham level during recovery (E) Loadings of Myristic acid (HMDB0000806) -> HNF4A interactions, one of the top interactions comprising factor 1 (D). As in (D), the interaction is strong in the sham, absent after AKI, and reappears during recovery after injury.

One of the challenges in the context of the metabolite estimation from transcriptomic data is that for a CCC event to happen the metabolites have to be secreted in the extracellular space, and sometimes transported inside of the receiver cell, to bind to the intended receptor (Supplementary Note 1). While ions and lipophilic substances can diffuse through membranes and vesicular transport systems exist, most molecules are transported via uni or bidirectional protein transporters. To account for this we collected information on importers and exporters from Recon3D^22^, Human Metabolic Atlas^23^, and TransportDB^47^ and calculated a weighted mean to assess their presence, keeping metabolites with net positive import or export estimates (Figure 4B, Methods).

To showcase the potential of MetalinksDB, we combined it with LIANA+ to infer CCC from spatial transcriptomics^26^ (Methods). Using LIANA+ we estimated spatially-weighted metabolite-receptor interactions for each spot across five slides of murine kidney, acquired before and following AKI (Figure 4C). We then used non-negative matrix factorization and identified three CCC patterns (factors), representing potential intercellular communication patterns (Figure 4C, Methods). All three factors, especially Factor 1 and Factor 2 showed a strong decrease of CCC events with disease onset and an increase during recovery (Figure 4D, S6). Most prominently, Factor 1 showed interactions of fatty acyls like linoleic, myristic, and dodecanoic acid with the hepatocyte nuclear factor 4 alpha (HNF4A) (Figure S7), which were disrupted in the early hours following AKI, and subsequently seen to be recovered in the late time points (Figure 4D,E; Figure S8). These CCC events are further characterized by colocalization of fatty acyl and mRNA in the medulla of the healthy and recovered kidney (Figure S8). Backing this discovery there is sufficient evidence that HNF4A binds to several different lipids as the ones observed (linoleic acid -> HNF4A, not it DB but suggested in the literature)^48^ and HNF4A signaling was found to drive recovery after AKI in mice^49^. Thus, here we show that combining MetalinksDB with LIANA+ facilitates the formulation of robust hypotheses of metabolite-mediated CCC relevant to specific disease contexts.

## 3. Discussion

In this paper, we report the assembly of a database called MetalinksDB, comprising the interactions between metabolite ligands and protein receptors as well as metabolic enzymes that produce and degrade these metabolites. Not only is it, to our knowledge, the most comprehensive database of its kind, but also the only one that is customizable through its flexible infrastructure. To make this database easily accessible, we built a web interface to allow users with no computational experience to customize, investigate, and download the database. We anticipate this to be relevant for the growing field of metabolite-mediated CCC, which will gain accuracy and impact with the advancements in single-cell metabolomics^50^.

We demonstrate the application of MetalinksDB in analyzing both bulk and spatial transcriptomics data. From bulk transcriptomic and metabolomic data, we find interactions and pathways deregulated in clear-cell renal cell carcinoma. We hypothesize that increased levels of ATP in connection with upregulated ATP exporters might lead to the activation of P2RY receptors that in turn could inhibit the sodium transporter ENaC. Additionally, we see downregulation of ENaC subunits (SCNN1_A,B,D,G_), a phenomenon that is associated with hyponatremia and metastasis in ccRCC ^41,42,46^. Additionally, we identified several disease-related factors in spatial transcriptomic data of murine AKI using the modular and flexible LIANA+ CCC framework. Interestingly, we found a HNF4A signature absent in disease states contributing mainly to the first disease-associated factors identified through NMF of local interactions. HNF4A was found to have a role in the AKI recovery of mice ^49^ and is known to bind to several lipids^48^, which makes it an interesting target to investigate further. Consequently, making use of diverse omics technologies, we demonstrate MetalinksDB’s utility not only in confirming interactions previously documented in the scientific literature but also in formulating hypotheses for novel ones.

The interactions described in the renal cancer use case were inferred using statistics representing metabolite or transcript deregulation in the bulk dataset. As such, they do not directly represent the deregulation of metabolite-protein binding. Similarly, the approach used to infer metabolite-receptor interactions from spatial transcriptomics is limited by inferring metabolite abundances from gene expression. This inference builds on the assumption that gene expression is a good measure of protein abundance and activity and later on that estimated enzyme abundance is a good measure of metabolite abundance. Thus, this approach ignores varying production/degradation rates between different proteins during metabolite production and the physicochemical properties of metabolites and their intra and extracellular surroundings. Thus, while here we infer putative metabolite-protein interactions, these remain only a hypothesis, to be validated.

As is commonly done in many biological databases, we had to decide which biological classes of protein-metabolite interactions to include and make a quality-coverage tradeoff, as putative interactions may originate from structural predictions or text mining, while the available manually curated interactions are limited to a few hundred. Users may want to choose via the web page or omnipath-client even cleaner yet smaller sets of interactions or investigate at a broad scale being more lenient to false positives. A further strategy to reduce false positives is to restrict the analysis to metabolites and receptors known to be present in specific biological contexts (such as tissues, niches, and diseases). Moreover, despite our efforts to enable flexible customization and to keep interactions of high confidence, we acknowledge that MetalinksDB contains interactions that may not reflect a direct molecular binding event, but rather a link that exists through close regulation of agents between metabolites and receptors. This is a consequence of how databases like STITCH, the current main source of MetalinksDB, are built. This highlights potential future directions such as the inclusion of other generalistic metabolite-protein databases ^54^. Moreover, experimental protocols that allow for a systematic characterization of the direct binding of protein and metabolites ^55,56^ will enable curation efforts in the future, for which MetalinkDB will be a suitable starting point.

Taken together, MetalinksDB provides a comprehensive and flexible resource for the growing field of metabolite-mediated CCC and will enhance data interpretation, particularly in studies where tissue context is of importance as shown in the examples of kidney diseases. Beyond the use of cell-cell communication, it will ease any metabolite-related enrichment task and enhance metabolic ^51,52^ - and prior knowledge-informed models^53^.

## 4. Methods

### 4.1 Knowledge Graph assembly

#### Interaction and annotation data

Information of metabolite protein-receptor interactions was obtained from STITCH, NeuronChat, and CellphoneDB. The detailed interactions and actions STITCH datasets were obtained from the webpage (http://stitch.embl.de/) after restricting the entries to human associations. The detailed interactions file as well as the actions file were then loaded and subset to interactions having an annotated mode. Following this, the remaining interactions were cut down to only have the desired modes of action (e.g. activation, binding, inhibition). The provided CIDs and Ensembl protein IDs were converted to HMDB and UniProt IDs using the pypath module of Omnipath^8^.

Directional information on which enzymes produce and degrade a metabolite was obtained from HMDB as well as GEMs ^20,22,23^. HMDB protein and metabolite information was downloaded as a .xml file from the HMDB webpage (https://hmdb.ca/downloads) and parsed to a data frame using xml.sax and xml.etree. HMDB reaction information was scraped using the request and BeautifulSoup package. The HMDBP IDs obtained from scraping were translated to UniProt IDs, using the mappings obtained from HMDB protein data. In later versions, the data is pulled from Omnipath, which follows a similar parsing strategy to obtain the data.

The GEM models were obtained via download as well (https://www.vmh.life/, https://github.com/SysBioChalmers/Human-GEM). For the metabolic enzyme resource, we transformed the models, consisting of a stoichiometric matrix and information about the genes and metabolites, into a data frame consisting of gene-metabolite associations and a directionality. Associations resulting from reactions annotated as reversible were assigned both directionality, except for proteins annotated as transporters. Missing identifiers were filled in by an ID translation table obtained from the HMDB metabolite data, as well as an in-house table (https://github.com/biocypher/metalinks) and a table obtained from the metaboliteIDmapping R package (https://github.com/yigbt/metaboliteIDmapping). Transporters were determined by the subsystem channel of the model and a direction was defined using the compartment annotation. Transport from an organelle to the cytosol as well as from the cytosol to the extracellular milieu was assigned as outwards and inwards vice versa. Both association lists were then combined and duplications were removed.

Further databases were leveraged to provide comprehensive annotation of metabolites and proteins. Uniprot data was downloaded via the API using the crossbar project (https://crossbar.kansil.org/project.php) BioCypher adapter. TransportDB2.0 and “Guide to Pharmacology” data were obtained from the respective web pages (http://www.membranetransport.org/transportDB2/index.html, https://www.guidetopharmacology.org/download.jsp).

The code handling the above tasks was forged into BioCypher adapters to allow proper versioning and transparency of assembly available at https://github.com/biocypher/metalinks. Executing the code yields several .csv files which can be loaded via Neo4j, which enables MetalinksDB to be queried using the cypher querying language ^57^.

During the knowledge graph assembly, several cutoffs are applied, which are summarized here for transparency. At first, we excluded all interactions from STITCH that had no annotated interaction mode or available HMDB and Uniprot identifiers after conversion. Secondly, we excluded all proteins that were not annotated as catalytic receptors, GPRCs, Ion channels or transporters in the “Guide to Pharmacology”. We further included only interaction that had an interaction mode of activation, inhibition or binding, for the latter we however excluded all interactions involving Ion channels or transporters, due to too complex biological interpretations. We applied a cutoff to the provided confidence level that comes with every STITCH connection, based on the distribution of manually curated interactions from CellphoneDB and NeuronChat (Figure S2). In brief, we investigated the STITCH confidence levels of interactions found in both the curated and STITCH data. The distribution gave us an impression that a substantial amount of true positives could be found, resulting in cutoffs of 300 for the database confidence score, 300 for the experiment score, 700 for the prediction score, and 900 for the combined one, the text mining was not taken into account. Finally, we excluded all metabolites that were not annotated as extracellular in the HMDB data.

This filtering strategy is a compromise from various priorities that other researchers may set differently. To address this problem we build a web interface, where these filtering cutoffs can be adjusted to the user’s interest.

### 4.2 Web interface

The web interface (https://metalinks.omnipathdb.org/) is based on the streamlit library that uses the neo4j driver to query data from the MetalinksDB knowledge graph. The code can be found here https://github.com/saezlab/metalinks_web. The graph interface is built on the drugst.one html infrastructure ^28^.

### 4.3 Data analysis

#### 4.3.1 Application on bulk transcriptomics and metabolomics from ccRCC Patient data

Differential abundance statistics for metabolomic and transcriptomic data were obtained from Dugourd et al. 2021 ^25^. As a resource for metabolite-mediated CCC interactions MetalinksDB was contextualized to metabolites found in kidney blood or urine and stronger cutoffs for the STITCH parameters were enforced to lower false positives (>500 Database, >500 Experiment and >700 Prediction). The remaining interactions were ranked by the mean of their t-values.

For the enrichment analysis, we downloaded the KEGG C2 set from MSigDB ^58^(https://zenodo.org/records/10200150) and metabolic pathway annotations from a metabolic ccRCC atlas ^30^ and created a univariate linear model for metabolites and transcripts on the connection level using decoupler^59^. The code for this calculation is available here (https://github.com/saezlab/metalinks/blob/main/nb/Figure_3.ipynb)

#### 4.3.2 Application on Kidney Injury model from Spatial Transcriptomics data

Five slides of AKI were obtained from a study by Dixon et al. ^60^. Spots were log normalized and genes were filtered for genes having more than 20 counts using the SCANPY package^61^. We inferred metabolite presence as described in LIANA+, and used it to infer local ligand-receptor communication events using spatially-weighted Cosine similarity^26^. Subsequently, we used a Gaussian radial kernel with a bandwidth of 100 to determine the spatial connectivities between spots. As a resource of metabolite-receptor interactions, we used a conservative set of connections mainly consisting of manually curated interactions. We set the negative metabolite values to 0 and calculated Cosine similarities between inferred metabolite presence and corresponding receptors/transporters per slide. We then concatenated all the local scores from all slides and performed a non-negative matrix factorization with 3 factors, as determined by the elbow method according to the LIANA+ defaults.

The graphical abstract as well as Figures 1, 3, 4, and S1 were created using Biorender. 20

## 5. Acknowledgements

C.S. was funded by the German Federal Ministry of Education and Research (Bundesministerium für Bildung und Forschung BMBF) MSCoreSys research initiative research core SMART-CARE (031L0212A). D.D. is supported by the European Union’s Horizon 2020 research and innovation program (860329 Marie-Curie ITN “STRATEGY-CKD”). D.T. was supported by the German Federal Ministry of Education and Research (BMBF) [031L0181B]; HPC/Exascale Centre of Excellence for Personalised Medicine in Europe [PerMedCoE; European Union Horizon 2020 program, grant no. 951773]. S.L. has received funding from the European Union’s Horizon 2020 research and innovation program (grant agreement No 965193 [DECIDER]).

We thank Pau Badia, Erick Armingol, Ruth Seurinck, Marlies Brouckaert, and all Saezlab members for helpful discussions.

## 7. Authors contributions

Conceptualization: AD, DD, JS

Data Curation: EF, AD, DD

Formal Analysis: EF, CS

Methodology: SL, AD, DD, DT, EF

Webpage: DT, EF

BioCypher backend: EF, SL

Visualization: EF

Funding acquisition: JS

Project administration: JS, AD, DD

Supervision: AD, DD, SL, JS

Writing – original draft: EF, DD, CS

Writing – review & editing: EF, DD, DT, SL, CS, AD, JS

## 8. References

1. Baker, S. A. & Rutter, J. Metabolites as signalling molecules. Nat. Rev. Mol. Cell Biol. 24, 355–374 (2023).

2. Haas, R. et al. Intermediates of metabolism: from bystanders to signalling molecules. Trends Biochem. Sci. 41, 460–471 (2016).

3. Armingol, E., Officer, A., Harismendy, O. & Lewis, N. E. Deciphering cell-cell interactions and communication from gene expression. Nat. Rev. Genet. 22, 71–88 (2021).

4. Dimitrov, D. et al. Comparison of methods and resources for cell-cell communication inference from single-cell RNA-Seq data. Nat. Commun. 13, 3224 (2022).

5. Efremova, M., Vento-Tormo, M., Teichmann, S. A. & Vento-Tormo, R. CellPhoneDB: inferring cell-cell communication from combined expression of multi-subunit ligand-receptor complexes. Nat. Protoc. 15, 1484–1506 (2020).

6. Jin, S. et al. Inference and analysis of cell-cell communication using CellChat. Nat. Commun. 12, 1088 (2021).

7. Noël, F. et al. Dissection of intercellular communication using the transcriptome-based framework ICELLNET. Nat. Commun. 12, 1089 (2021).

8. Türei, D. et al. Integrated intra- and intercellular signaling knowledge for multicellular omics analysis. Mol. Syst. Biol. 17, (2021).

9. Husted, A. S., Trauelsen, M., Rudenko, O., Hjorth, S. A. & Schwartz, T. W. GPCR-Mediated Signaling of Metabolites. Cell Metab. 25, 777–796 (2017).

10. Chantranupong, L., Wolfson, R. L. & Sabatini, D. M. Nutrient-sensing mechanisms across evolution. Cell 161, 67–83 (2015).

11. Wolfson, R. L. & Sabatini, D. M. The Dawn of the Age of Amino Acid Sensors for the mTORC1 Pathway. Cell Metab. 26, 301–309 (2017).

12. Lawrence, R. E. & Zoncu, R. The lysosome as a cellular centre for signalling, metabolism and quality control. Nat. Cell Biol. 21, 133–142 (2019).

13. Zhao, W., Johnston, K. G., Ren, H., Xu, X. & Nie, Q. Inferring neuron-neuron communications from single-cell transcriptomics through NeuronChat. Nat. Commun. 14, 1128 (2023).

14. Garcia-Alonso, L. et al. Single-cell roadmap of human gonadal development. Nature 607, 540–547 (2022).

15. Vento-Tormo, R. et al. Single-cell reconstruction of the early maternal-fetal interface in humans. Nature 563, 347–353 (2018).

16. Zhang, Y. et al. Cellinker: a platform of ligand-receptor interactions for intercellular communication analysis. Bioinformatics (2021) doi:10.1093/bioinformatics/btab036.

17. Jakobsson, J. E. T., Spjuth, O. & Lagerström, M. C. scConnect: a method for exploratory analysis of cell-cell communication based on single-cell RNA-sequencing data. Bioinformatics 37, 3501–3508 (2021).

18. Harding, S. D. et al. The IUPHAR/BPS guide to PHARMACOLOGY in 2022: curating pharmacology for COVID-19, malaria and antibacterials. Nucleic Acids Res. 50, D1282–D1294 (2022).

19. Zheng, R. et al. MEBOCOST: Metabolic Cell-Cell Communication Modeling by Single Cell Transcriptome. BioRxiv (2022) doi:10.1101/2022.05.30.494067.

20. Wishart, D. S. et al. HMDB 5.0: the human metabolome database for 2022. Nucleic Acids Res. 50, D622–D631 (2022).

21. Szklarczyk, D. et al. STITCH 5: augmenting protein-chemical interaction networks with tissue and affinity data. Nucleic Acids Res. 44, D380–4 (2016).

22. Brunk, E. et al. Recon3D enables a three-dimensional view of gene variation in human metabolism. Nat. Biotechnol. 36, 272–281 (2018).

23. Robinson, J. L. et al. An atlas of human metabolism. Sci. Signal. 13, (2020).

24. Lobentanzer, S. et al. Democratizing knowledge representation with BioCypher. Nat. Biotechnol. 41, 1056–1059 (2023).

25. Dugourd, A. et al. Causal integration of multi-omics data with prior knowledge to generate mechanistic hypotheses. Mol. Syst. Biol. 17, e9730 (2021).

26. Dimitrov, D. et al. LIANA+: an all-in-one cell-cell communication framework. BioRxiv (2023) doi:10.1101/2023.08.19.553863.

27. Lobentanzer, S., et al. Democratising Knowledge Representation with BioCypher. arXiv (2022) doi:10.48550/arxiv.2212.13543.

28. Maier, A. et al. Drugst.One - A plug-and-play solution for online systems medicine and network-based drug repurposing. arXiv (2023) doi:10.48550/arxiv.2305.15453.

29. Linehan, W. M., Srinivasan, R. & Schmidt, L. S. The genetic basis of kidney cancer: a metabolic disease. Nat. Rev. Urol. 7, 277–285 (2010).

30. Hakimi, A. A. et al. An integrated metabolic atlas of clear cell renal cell carcinoma. Cancer Cell 29, 104–116 (2016).

31. Shuch, B., Linehan, W. M. & Srinivasan, R. Aerobic glycolysis: a novel target in kidney cancer. Expert Rev. Anticancer Ther. 13, 711–719 (2013).

32. Solini, A., Usuelli, V. & Fiorina, P. The dark side of extracellular ATP in kidney diseases. J. Am. Soc. Nephrol. 26, 1007–1016 (2015).

33. Menzies, R. I., Tam, F. W., Unwin, R. J. & Bailey, M. A. Purinergic signaling in kidney disease. Kidney Int. 91, 315–323 (2017).

34. Dwyer, K. M., Kishore, B. K. & Robson, S. C. Conversion of extracellular ATP into adenosine: a master switch in renal health and disease. Nat. Rev. Nephrol. 16, 509–524 (2020).

35. Kanehisa, M., Furumichi, M., Tanabe, M., Sato, Y. & Morishima, K. KEGG: new perspectives on genomes, pathways, diseases and drugs. Nucleic Acids Res. 45, D353–D361 (2017).

36. Tsilosani, A., Gao, C. & Zhang, W. Aldosterone-Regulated Sodium Transport and Blood Pressure. Front. Physiol. 13, 770375 (2022).

37. Anders, H.-J., Banas, B. & Schlöndorff, D. Signaling danger: toll-like receptors and their potential roles in kidney disease. J. Am. Soc. Nephrol. 15, 854–867 (2004).

38. Chen, H. et al. Combined clinical phenotype and lipidomic analysis reveals the impact of chronic kidney disease on lipid metabolism. J. Proteome Res. 16, 1566–1578 (2017).

39. Lucarelli, G. et al. Integration of lipidomics and transcriptomics reveals reprogramming of the lipid metabolism and composition in clear cell renal cell carcinoma. Metabolites 10, (2020).

40. Hanukoglu, I. & Hanukoglu, A. Epithelial sodium channel (ENaC) family: Phylogeny, structure-function, tissue distribution, and associated inherited diseases. Gene 579, 95–132 (2016).

41. Zheng, Q. et al. Inactivation of epithelial sodium ion channel molecules serves as effective diagnostic biomarkers in clear cell renal cell carcinoma. Genes Genomics 45, 855–866 (2023).

42. Nobel, Y. R. et al. Pseudohypoaldosteronism type 1 due to novel variants of SCNN1B gene. Endocrinol. Diabetes Metab. Case Rep. 2016, 150104 (2016).

43. Kaur, J. & Dora, S. Purinergic signaling: Diverse effects and therapeutic potential in cancer. Front. Oncol. 13, 1058371 (2023).

44. Corriden, R. & Insel, P. A. Basal release of ATP: an autocrine-paracrine mechanism for cell regulation. Sci. Signal. 3, re1 (2010).

45. Schachter, J. B., Li, Q., Boyer, J. L., Nicholas, R. A. & Harden, T. K. Second messenger cascade specificity and pharmacological selectivity of the human P2Y1-purinoceptor. Br. J. Pharmacol. 118, 167–173 (1996).

46. Jeppesen, A. N., Jensen, H. K., Donskov, F., Marcussen, N. & von der Maase, H. Hyponatremia as a prognostic and predictive factor in metastatic renal cell carcinoma. Br. J. Cancer 102, 867–872 (2010).

47. Elbourne, L. D. H., Tetu, S. G., Hassan, K. A. & Paulsen, I. T. TransportDB 2.0: a database for exploring membrane transporters in sequenced genomes from all domains of life. Nucleic Acids Res. 45, D320–D324 (2017).

48. Wisely, G. B. et al. Hepatocyte nuclear factor 4 is a transcription factor that constitutively binds fatty acids. Structure 10, 1225–1234 (2002).

49. Kirita, Y., Wu, H., Uchimura, K., Wilson, P. C. & Humphreys, B. D. Cell profiling of mouse acute kidney injury reveals conserved cellular responses to injury. Proc Natl Acad Sci USA 117, 15874–15883 (2020).

50. Seydel, C. Single-cell metabolomics hits its stride. Nat. Methods 18, 1452–1456 (2021).

51. Wagner, A. et al. Metabolic modeling of single Th17 cells reveals regulators of autoimmunity. Cell 184, 4168–4185.e21 (2021).

52. Alghamdi, N. et al. A graph neural network model to estimate cell-wise metabolic flux using single-cell RNA-seq data. Genome Res. 31, 1867–1884 (2021).

53. Lotfollahi, M. et al. Biologically informed deep learning to query gene programs in single-cell atlases. Nat. Cell Biol. 25, 337–350 (2023).

54. Zhao, T. et al. Prediction and collection of protein-metabolite interactions. Brief. Bioinformatics 22, (2021).

55. Piazza, I. et al. A Map of Protein-Metabolite Interactions Reveals Principles of Chemical Communication. Cell 172, 358–372.e23 (2018).

56. Hicks, K. G. et al. Protein-metabolite interactomics of carbohydrate metabolism reveal regulation of lactate dehydrogenase. Science 379, 996–1003 (2023).

57. Francis, N. et al. Cypher: an evolving query language for property graphs. in Proceedings of the 2018 International Conference on Management of Data - SIGMOD ’18 1433–1445 (ACM Press, 2018). doi:10.1145/3183713.3190657.

58. Liberzon, A. et al. Molecular signatures database (MSigDB) 3.0. Bioinformatics 27, 1739–1740 (2011).

59. Badia-I-Mompel, P. et al. decoupleR: ensemble of computational methods to infer biological activities from omics data. Bioinformatics Advances 2, vbac016 (2022).

60. Dixon, E. E., Wu, H., Muto, Y., Wilson, P. C. & Humphreys, B. D. Spatially resolved transcriptomic analysis of acute kidney injury in a female murine model. J. Am. Soc. Nephrol. 33, 279–289 (2022).

61. Wolf, F. A., Angerer, P. & Theis, F. J. SCANPY: large-scale single-cell gene expression data analysis. Genome Biol. 19, 15 (2018).

62. Wang, Y.-P. & Lei, Q.-Y. Metabolite sensing and signaling in cell metabolism. Signal Transduct. Target. Ther. 3, 30 (2018).

63. Alberts, B., et al. *Molecular Biology of the Cell*. Sixth Edition. Q. Rev. Biol. 90, 343–343 (2015).

64. Gu, C., Kim, G. B., Kim, W. J., Kim, H. U. & Lee, S. Y. Current status and applications of genome-scale metabolic models. Genome Biol. 20, 121 (2019).

65. Djoumbou-Feunang, Y. et al. BioTransformer: a comprehensive computational tool for small molecule metabolism prediction and metabolite identification. J. Cheminform. 11, 2 (2019).

66. Bansal, P. et al. Rhea, the reaction knowledgebase in 2022. Nucleic Acids Res. 50, D693–D700 (2022).

67. Gillespie, M. et al. The reactome pathway knowledgebase 2022. Nucleic Acids Res. 50, D687–D692 (2022).

