## Supplementary Information for "MetalinksDB: a flexible and contextualizable resource of metabolite-protein interactions"

### Supplementary Materials

#### Supplementary Files

Supplementary Table 1: Databases for metabolite-mediated CCC

| Name | Size [Interactions] | Other contents | Curation | Availability | Source |
|---|---|---|---|---|---|
| **scConnect** | 645 | protein - protein interactions | Database | Open | GtP |
| **Cellinker** | 254 | protein - protein interactions | Databases | Open | GtP |
| **NeuronChat** | <100 | protein - protein interactions | Manual | Open | |
| **CellphoneDB** | ~240 | protein - protein interactions | Manual | Open | |
| **MEBOCOST** | ~421 | **-** | Databases, Text-mining | Restricted* | |

*Supplementary File 1: Metabolic signaling*

In the field of nutrient sensing and metabolite signaling, multiple sometimes conflicting concepts of metabolic signaling exist [1,9,62]. This can impact the computation of metabolite-mediated CCC, since the concept influences what interactions to include in the prior knowledge. The diversity within the concepts may originate from the fact that most proteins can't surpass cell membranes, while metabolites often do. They can because of their small size and chemical nature, which allows them to either diffuse through the membrane or use passive or active transport through the cell membranes [63]. Inside the cell, metabolites can interact with proteins in several ways, either as substrates to metabolic enzymes, allosteric regulators or scaffold molecules for protein complexes[55]. The allosteric regulation includes activation of proteins as in the case of nuclear receptors or inhibition due to binding to reactive sites.

For a cell, the consequences of a metabolite-protein interaction can range from very strong consequences, due to the activation of a signaling cascade in case of a binding event to a membrane receptor or a change in transcription due to binding to a nuclear receptor, to very weak consequences that may be the result of the metabolizing event of a sugar or lipid that happens hundreds of times per second. However, many weak interaction effects can result in a strong cellular effect as well, most often through the signaling capacity of accumulating upstream or downstream metabolites [1,62]. Since the cellular effect is most often the parameter of interest it can therefore be challenging to classify the interactions into useful categories. These categories would however allow us to clearly define which interactions to include.

In our eyes, a classification of signaling interactions needs to give information on I) if the signaling should be focussed on the signal amplification strength of the first messenger II) the downstream distance where signal amplification is allowed to happen, and III) the strength or kind of morphological change of the cell after signal integration.

Consequently, three natural ways of classifying all interactions of an extracellular metabolite with a target protein of a cell emerge. Classifying by strength or kind of morphological change of a cell may be of interest for researchers using techniques mostly focussed on morphology, for example, microscopists or physicians. Due to its apathy towards the underlying mechanisms of the change, it may be unsuited for mechanistic approaches as we try to investigate here.

The classification focusing on where in the process the signal is integrated in a way that it can be amplified is interesting for people focussed on intracellular pathways and networks and may be of interest to metabolic CCC if the methods are more mature. So far it is hard to estimate if there will be a useful connection between certain groups of first messengers and downstream amplification mechanisms.

The last option seems best suited for metabolic CCC since it focuses on the first messenger. It separates the different signaling modes from their downstream cascade and classifies them into the signal amplification capability of the direct target. As can be seen in Figure S1, as best examples for a clear first messenger-signal amplification relationship A and B the signaling molecule binds to an outer membrane - or nuclear receptor. These receptors often have a clear downstream effect that already amplifies the signal, even if further signal integration may happen.

Similar to this, C describes first messengers that bind to ion channels or transporters that allow an in - or efflux of molecules that may have signaling properties (Figure S1). A more subtle interaction with a receiver cell is any interaction with an enzyme inside a cell that affects the enzyme's function (Figure S1 E). Finally, there are two classes of signaling interactions in which there is no defined metabolite-protein interaction, that D the contact-dependent exchange of molecules between two cells, and F the change of a metabolic rate through the higher or lower abundance of a metabolite.

This definition of latter cases as signaling is of further relevance, when considering to include the results of several interactomics screens into the metabolite-receptor interactions list [55,56]. Since in their screens, a protein-metabolite interaction can stand for allosteric regulation of the protein, but also interactions as the substrate of the enzyme, we have to decide if we include the signaling events of the accumulation of upstream or downstream product of a reaction into account. In this manuscript, we decided to leave these cases out and therefore not include the interactomics screen results unless the interaction's nature is classified.

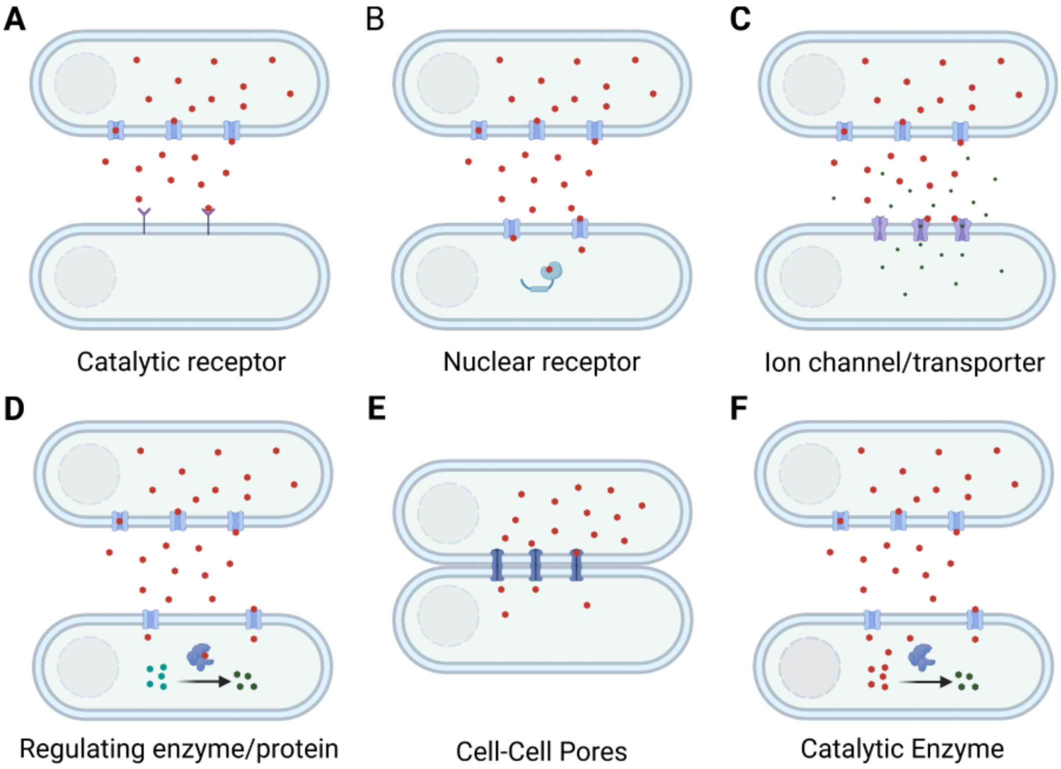


***Figure S1: Modes of metabolite-mediated CCC***
Metabolite-mediated CCC can be classified in several communication types. While some of the receptors are located in the membrane (A)(C), intracellular receptors exist as well (B)(D)(F). (E) is a special case of signaling involving direct contact of cells through channels. All modes differ in the way molecule binding or exchange is amplified.

Another advantage of this strategy is that we can focus on certain cases, that for example incorporate a certain type of receptor and can make specific assumptions that ease the modeling for these cases. For instance in case of A, we don't need to think about importing transporters, while for B,C,E,F we have to. Similarly, different prior knowledge resources are needed for the different cases that may change the structure of algorithms.

*Supplementary File 2: Discussion of databases used*

The main source of information of the ligand-receptor database is the STITCH database which serves as an extensive resource for over 20 million protein interactions, incorporating a diverse range of information from experimental data, databases, computational predictions, and text mining[20]. This comprehensive database provides crucial insights into the mode of binding, such as activation, inhibition, or binding, along with confidence scores for individual sources and combined analyses. Notably, the mode of binding is highly significant for downstream analysis; however, our understanding of what a binding event truly entails is mostly limited to activation events.

Despite the assigned confidence levels, determining the validity of a connection can be challenging, especially when limited information is available. Thus, users must establish their own context-specific cutoffs based on relevant ground truth data. Neo4j, with its interactive graph representations and Cypher query language, presents an ideal tool for contextualizing protein interactions[57]. For this purpose, information from various secondary sources is utilized, predominantly HMDB, Recon3D, and UniProt, to obtain attributes of protein, metabolite, and association nodes. The validity of these annotations for diseases, pathways or tissue context is also under debate as previous investigations showed unsatisfying accuracy of annotations like the cellular location of proteins[4].

Our production degradation resource is primarily based on the Recon3D metabolic model and the reaction webpages of the Human Metabolome Database (HMDB). Despite identifying some inaccuracies in the interactions, the Recon3D model remains widely accepted within the metabolomics field[64]. Each reaction on the Recon3D webpage is supported by evidence from the literature, enabling users to verify annotations. Another available model of human metabolism is the human HMR model, which has better curated fatty acid metabolism [23,64].

HMDB sources most of its reactions from KEGG, another pathway database, though the latter's lack of open access presents challenges for direct usage[35]. Additionally, a significant portion of HMDB interactions are derived from BioTransformer, a model for predicting metabolism, such as lipid breakdown[65]. However, we approached these interactions with caution and frequently excluded them from our analysis.

Alternative databases, such as Rhea or Reactome, store metabolite-enzyme interactions but possess limitations due to the absence of directionality assignment or protein assignment for reactions, rendering them less useful for our purposes [66,67]. If there were a high user demand to include reactions from these databases, it would be possible to do so, potentially through the use of BioCypher adapters. This would allow users with different requirements or preferences, such as those unconcerned with reaction directionality, to incorporate additional associations into their analyses.

An aspect that should be further investigated is the impact of set size on the accuracy of the metabolite estimation. So far, tools like Cellphone and NeuronChat use one and three enzymes as sets, while MEBOCOST and MetalinksDB use sizes from 1 to over 50 enzymes in a set. An approach to restrict the impact here would be to use the geometric mean or maximum value, that would correct for or be agnostic to the number of enzymes in a set.

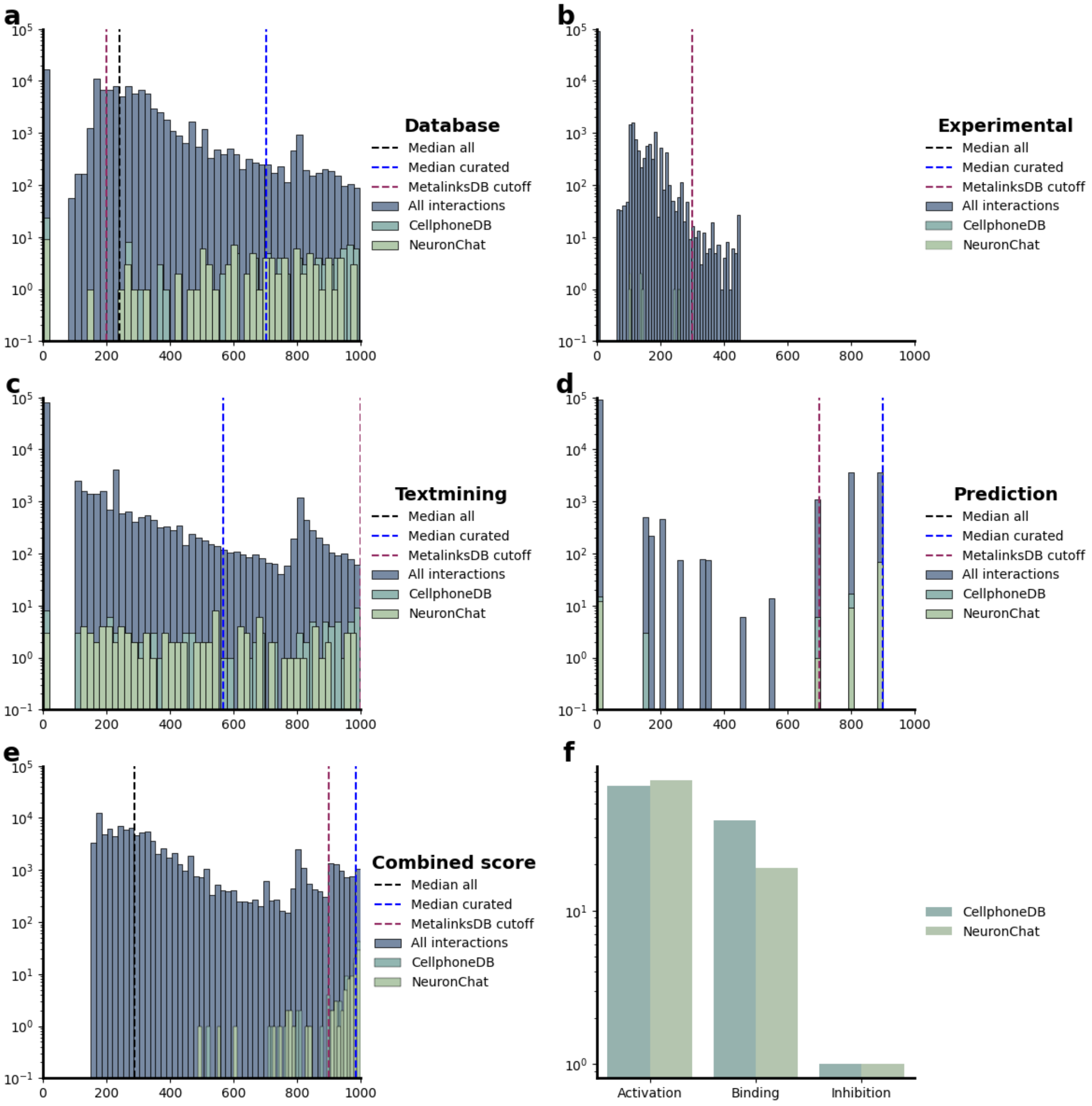


**Supplementary Figure S*2:* Comparison of STITCH confidence scores with manual curation**

(a)-(e) Database, Experimental, Textmining, Textmining, and Combined confidence scores of MetalinksDB interactions. Interactions that are also in manually curated databases (CellphoneDB, NeuronChat) are shown in green. Manually curated interactions have high Database, Prediction, and Combined scores, while few have experimental values and text mining appears evenly distributed. (f) Histogram of interaction classification of CellphoneDB and NeuronChat, showing that most of their interactions are activating or binding.

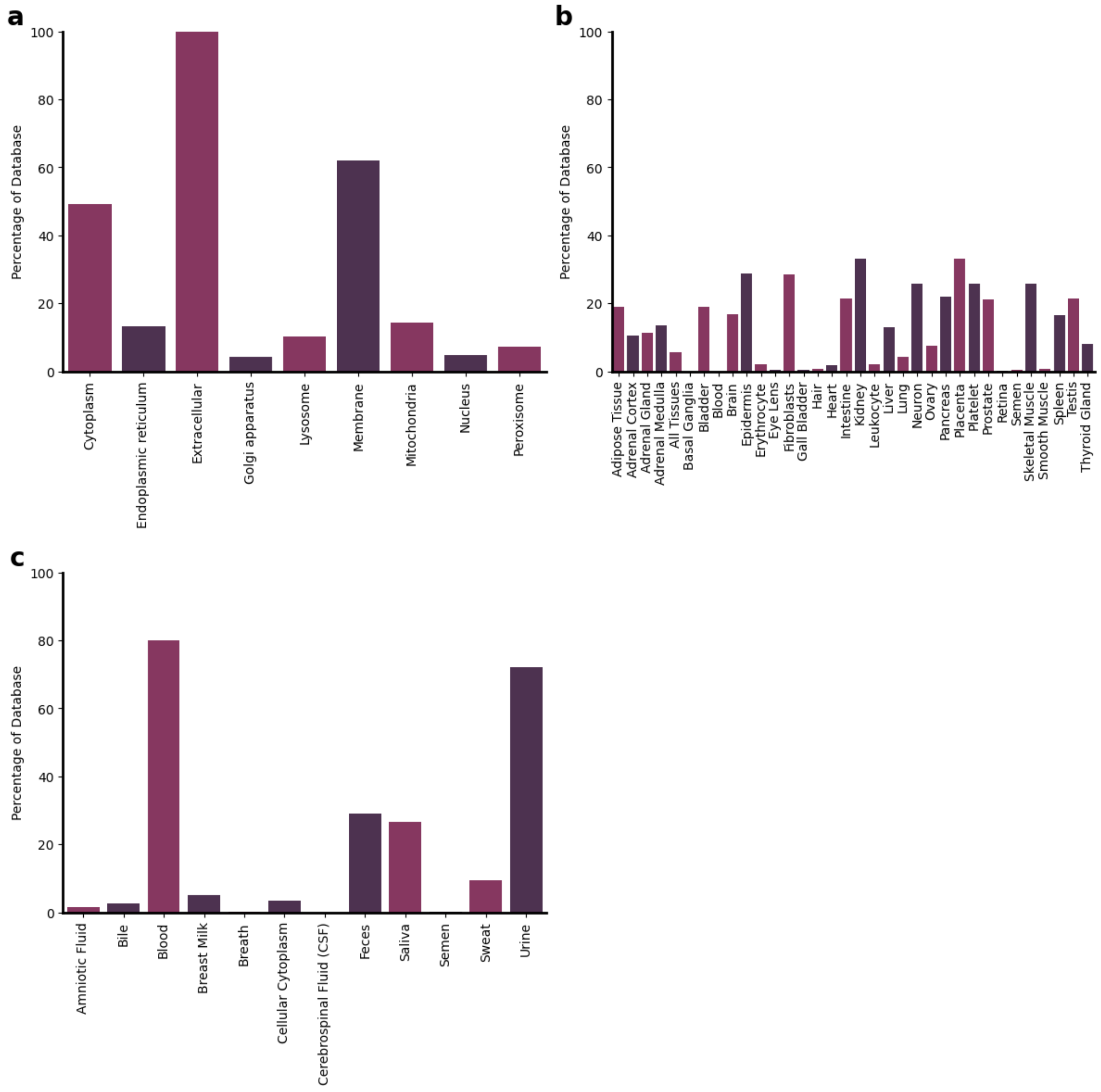


**Supplementary Figure S3: Fractions of metabolites in MetalinksDB belonging to annotation classes**

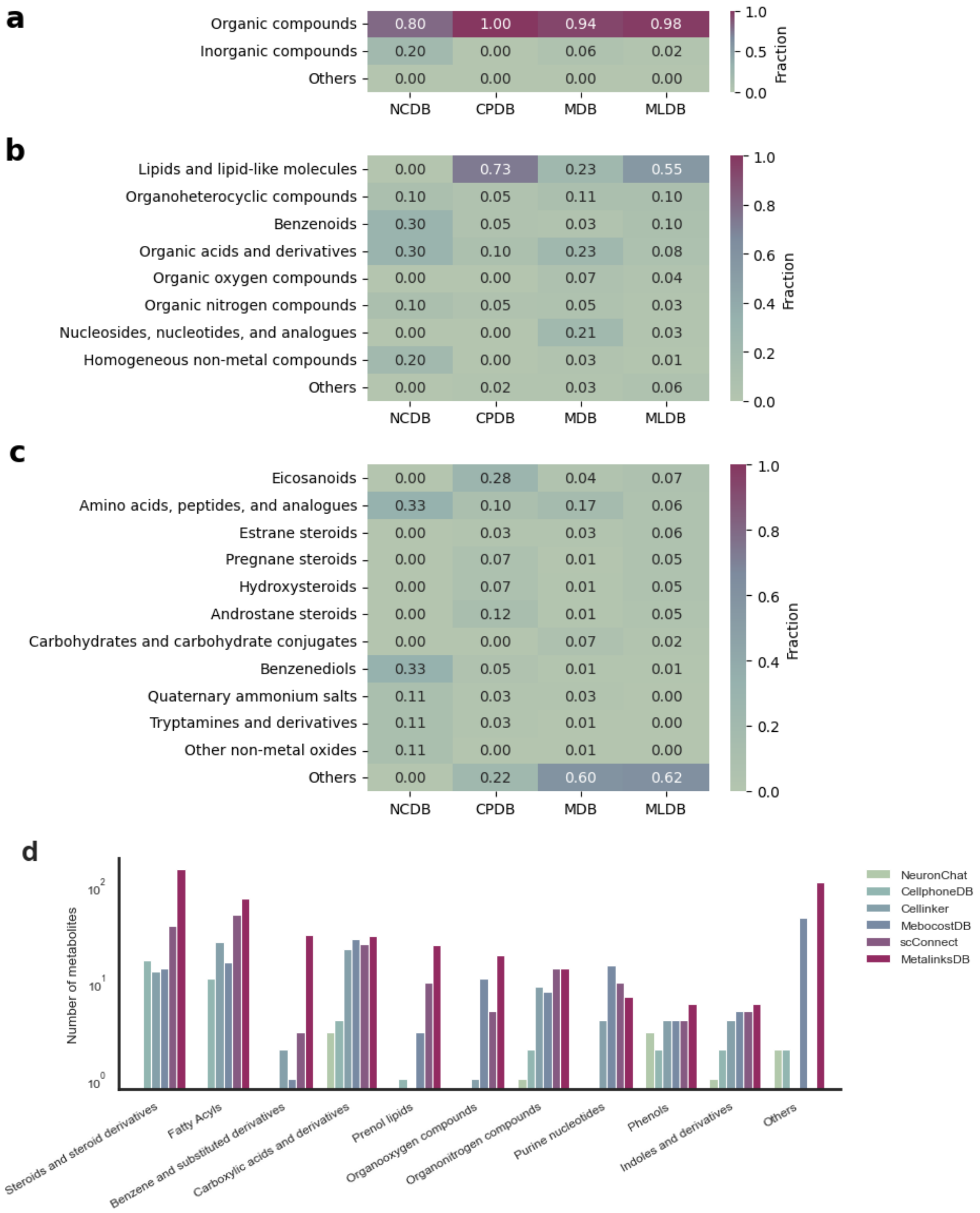


**Supplementary Figure S4: Metabolite classes through the database**

(A)-(C) Heatmaps of fractions of metabolite classes, comparable to Figure 2B. (D) Histogram of absolute values of metabolite classes.

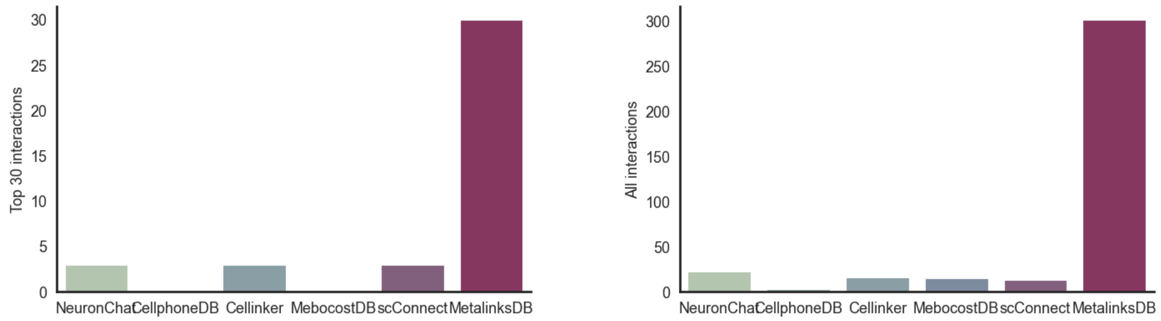


**Supplementary Figure S5: MetalinksDB Kidney analysis interactions found in other databases**

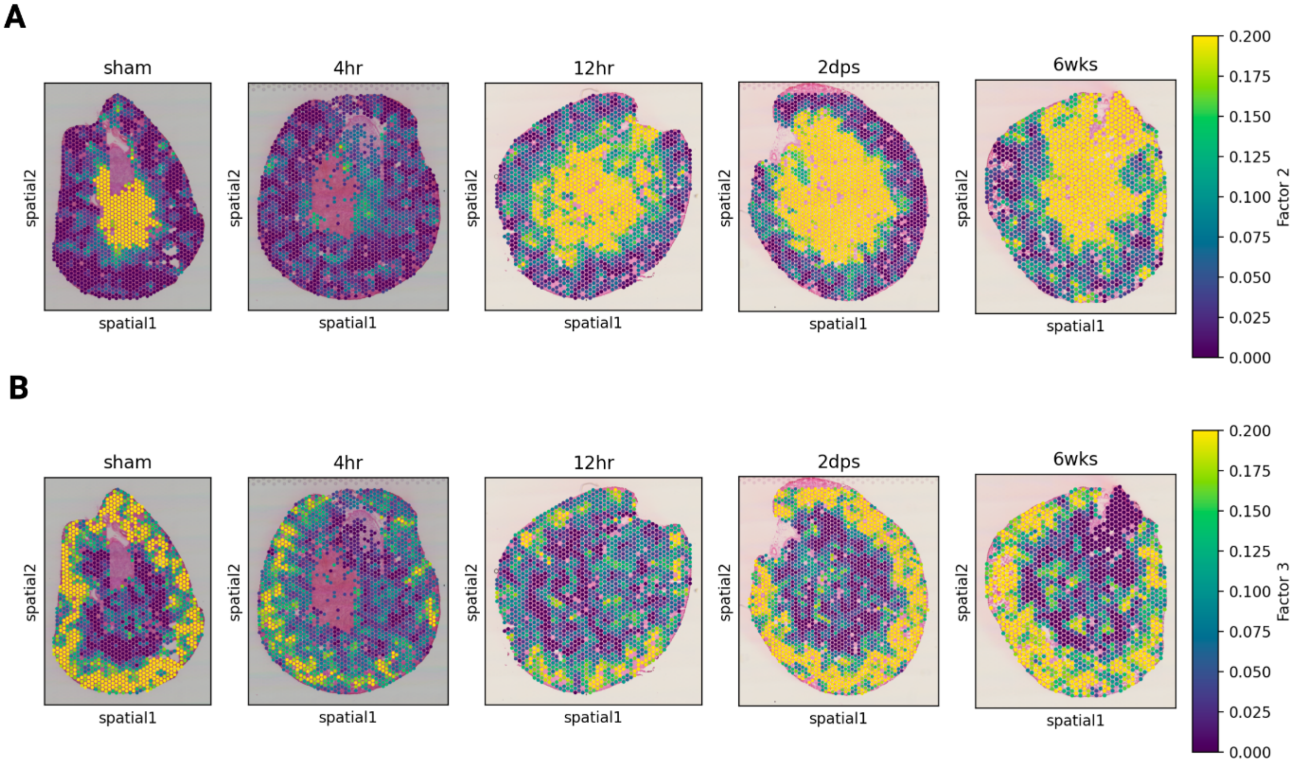


**Supplementary Figure S6: Spot scores of factors 2 and 3 from AKI factor analysis**

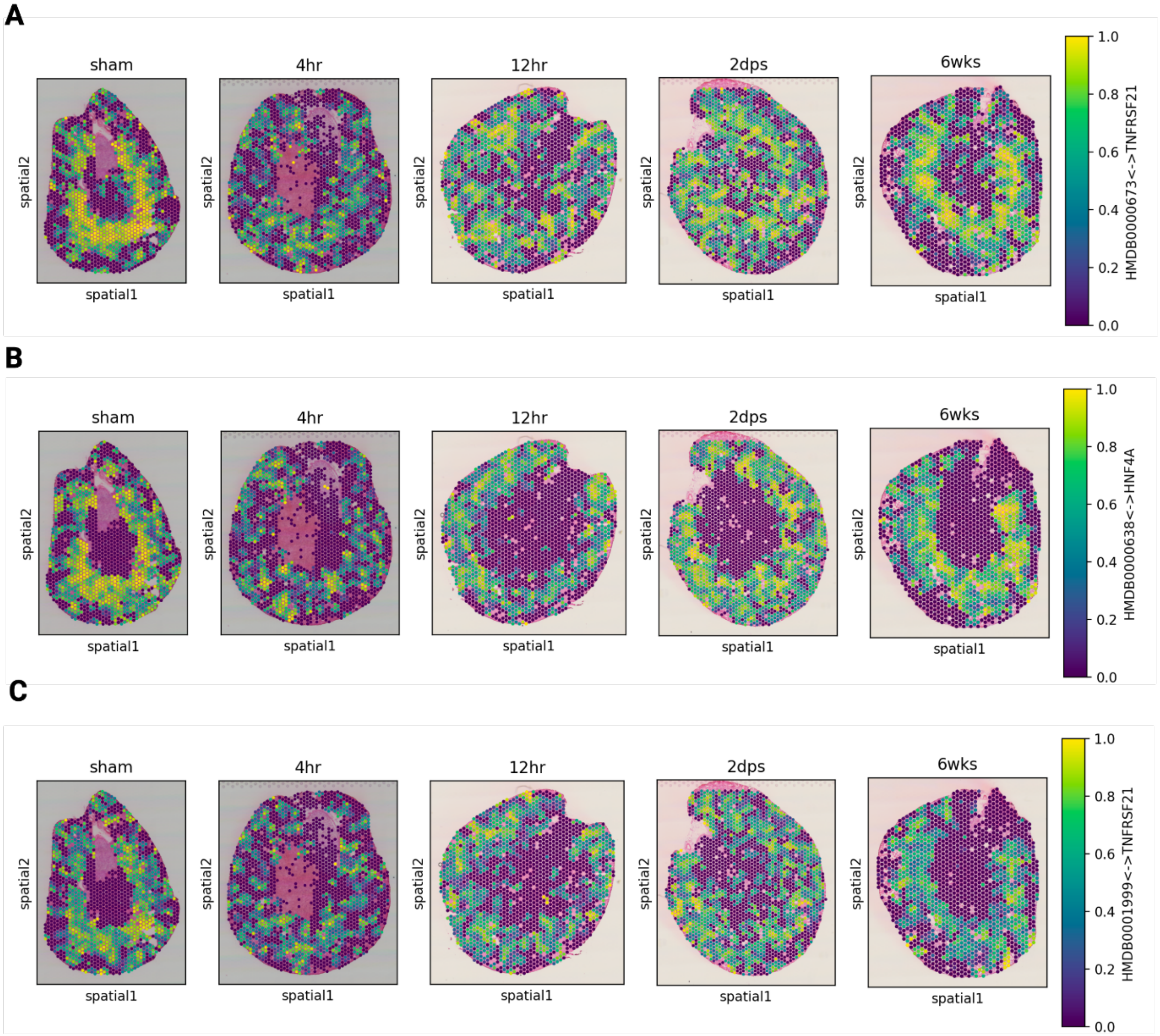


**Supplementary Figure S7: Spot scores of top interactions of Factor 1**

(A) Interaction scores of Linoleic acid (HMDB0000673) with tumor necrosis factor receptor superfamily member 21 (TNFRSF21) (B) Interaction scores of Dodecanoic acid (HMDB0000638) with HNF4A (C) Interaction scores of Eicosapentaenoic acid (HMDB0001999) with TNFRSF21

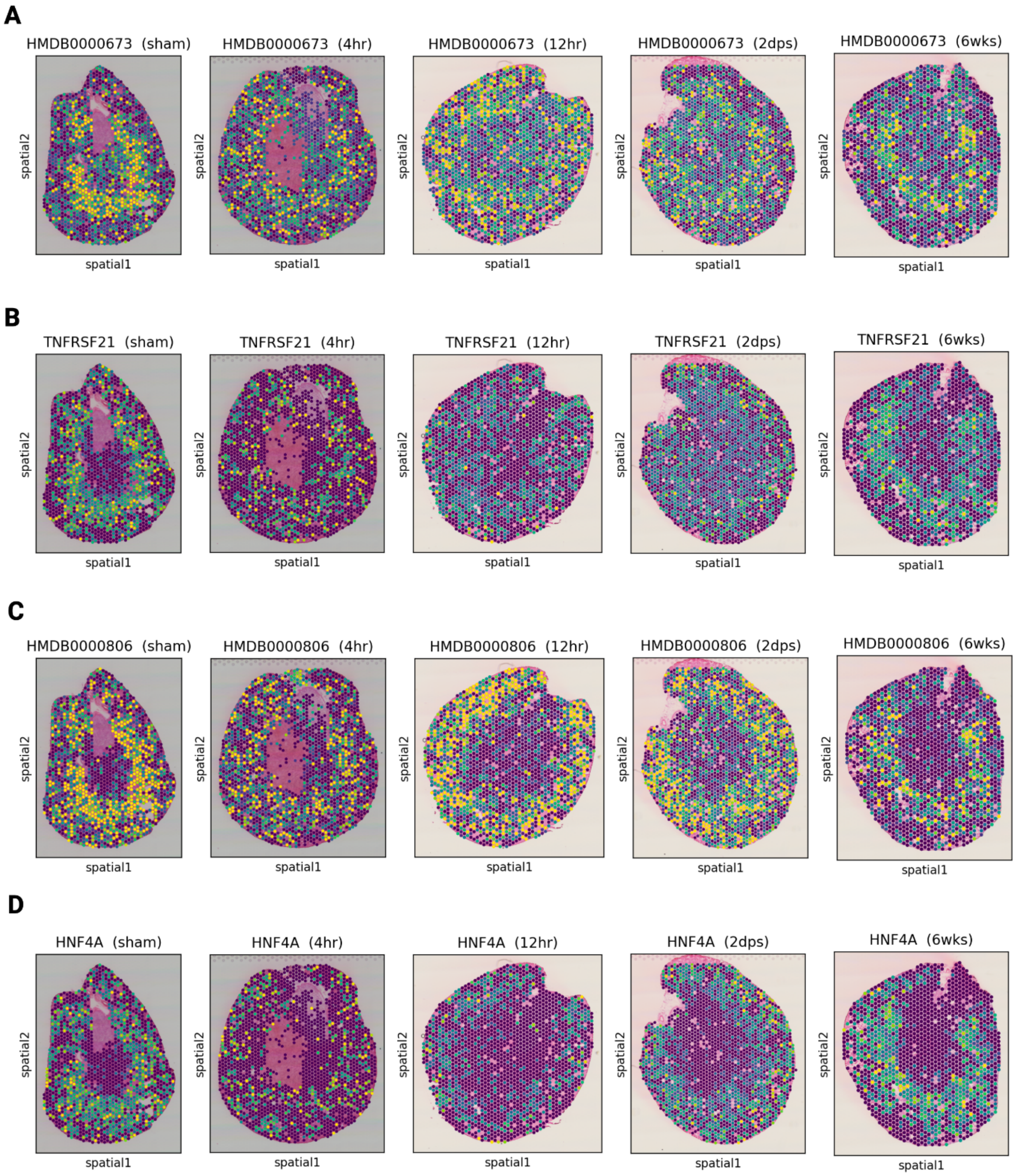


**Supplementary Figure S8: Spatial estimations of Acids and receptors involved**

(A) Abundance estimation of linoleic acid (HMDB0000673) (B) Expression of TNFRSF21 (C) Abundance estimation of myristic acid (HMDB0000806) (D) Expression of HNF4A
